# SENSORY MAPS IN THE TELENCEPHALIC PALLIUM OF GOLDFISH

**DOI:** 10.64898/2026.03.25.714251

**Authors:** F.M. Ocaña, A. Gómez, C. Salas, F. Rodríguez

## Abstract

The functional organization of the teleost telencephalic pallium remains poorly understood, particularly regarding the presence of modality-specific sensory domains and their topographic arrangement. Here, we used in vivo wide-field voltage-sensitive dye imaging to map sensory-evoked neural activity across the dorsal surface of the telencephalic pallium of adult goldfish. Somatosensory, auditory, gustatory, and visual stimulation revealed distinct, modality-specific domains primarily located within the dorsomedial (Dm) and dorsolateral (Dl) pallium. Within Dm, somatosensory and auditory stimuli activated partially overlapping territories in the caudal subregion (Dm4), exhibiting clear somatotopic and tonotopic organization along the mediolateral axis. Gustatory stimulation selectively engaged Dm3, where different tastants activated spatially distinct but partially overlapping domains. A more rostral subregion (Dm2) responded only to high-intensity somatosensory stimulation, suggesting involvement in processing negatively valenced inputs. Visual stimulation activated a circumscribed area within the dorsolateral pallium (Dld2),that closely matched cytoarchitectural boundaries. Pharmacological blockade of ionotropic glutamate receptors markedly reduced sensory-evoked responses, indicating that these maps depend on glutamatergic synaptic transmission. Together, these findings show that the goldfish pallium contains distinct, spatially organized sensory representations and a refined internal functional architecture. This organization suggests that pallial topographic sensory maps may not be exclusive to mammals and birds. Based on these results, we propose that dorsomedial and dorsolateral pallial regions may be functionally comparable to components of the mammalian mesocortical network, more than to the pallial amygdala or the neocortex. This framework provides a new perspective on pallial organization in teleosts and contributes to understanding the evolutionary origins of the vertebrate pallium.

**HIGHLIGHTS:**

- Voltage-sensitive dye imaging was used to map sensory responses in the goldfish pallium.
- Distinct sensory areas for somatosensory, auditory, gustatory, and visual modalities were identified.
- Some sensory regions in Dm show topographically organized maps.
- Functional segregation suggests a complex, non-diffuse pallial organization.
- Findings support a novel hypothesis linking Dm and Dld to mammalian mesocortical regions.

## INTRODUCTION

The telencephalic pallium is one of the most anatomically diverse structures in the vertebrate brain (Nieuwhenhuys et al, 1998; Striedter & Northcutt, 2020; Wulliman & Vernier, 2009). In actinopterygians (ray-finned fishes), its architecture differs markedly from that of other groups primarily due to a unique developmental process known as eversion, in which the telencephalic walls fold outward rather than inward. This contrasts with the evagination seen in sarcopterygians (lobe-finned fishes and tetrapods), and likely results in a rearrangement of pallial topography (Folgueira & Clarke, 2024; Nieuwenhuys, 2011). In addition, particularly within teleosts, the pallium exhibits a high degree of cytoarchitectural differentiation and regional specialization, further complicating attempts to establish homologies with pallial subdivisions of land vertebrates. Moreover, its functional organization remains incompletely understood.

While there is broad consensus that teleosts possess homologues of the olfactory pallium, hippocampus, and pallial amygdala, the presence of a region comparable to the neocortex of mammals remains highly controversial (Butler, 2000; Mueller et al., 2011; Striedter & Northcutt, 2021; Wullimann & Mueller, 2004; Yamamoto et al 2007). Anatomically, the teleost pallium is commonly divided into medial (Dm) and lateral (Dl) regions, separated by the sulcus ypsiloniformis. Dl, especially its ventral part (Dlv), is widely considered homologous to the hippocampus, based on convergent developmental, molecular, and functional evidence, including its well-established role in spatial navigation and memory (Furlan et al 2017; Ganz et al., 2014; Hegarty et al., 2024; Rodríguez et al., 2002). The Dm region, in contrast, is widely regarded as homologous to the pallial amygdala of other vertebrates on the basis of its topological position, molecular signatures, gene expression profile, connectivity with subpallial and hypothalamic regions, and its role in emotional processing (Lal et al 2018; Northcutt, 2006; Portavella et al., 2004; Porter & Mueller, 2020).

However, other hypotheses propose that sensory-recipient zones within both Dl and Dm could be comparable to neocortical areas in mammals (Mueller et al 2011; Striedter & Northcutt, 2021; Yamamoto et al 2007). This view arises from the observation that these pallial regions receive inputs from multiple sensory modalities—including auditory, visual, gustatory, and somatosensory—relayed through diencephalic relay centers (Saidel et al., 2001; Yamamoto & Ito, 2008). Nonetheless, unlike in tetrapods where sensory inputs are relayed through the dorsal thalamus, teleosts receive them mainly from the preglomerular complex, a highly derived structure originating from the midbrain and posterior diencephalon, whose homology to the thalamus remains unresolved (Bloch et al 2020; Ishikawa et al 2007; Northcutt, 2008; Yamamoto & Ito, 2005). Furthermore, it is not resolved whether the teleost pallium hosts discrete, modality-specific domains organized into spatially ordered maps, akin to those of the neocortex, or whether it functions instead as a more integrative, limbic-like multimodal domain.

A central question in this debate, therefore, is whether the teleost pallium contains discrete, unimodal sensory domains organized into spatially ordered maps—such as somatotopic, tonotopic, or retinotopic representations—or whether it lacks such topographic organization. The prevailing view, shaped by longstanding assumptions in comparative neuroanatomy, holds that the presence of spatially ordered sensory maps is a defining feature of advanced pallial architectures that are unique to “higher vertebrates” -i.e., mammals and birds- and absent in fishes (e.g., Damasio & Carvalho, 2013; Feinberg & Mallatt, 2016; Graziano, 2019; LeDoux, 2021). This assumption has often led to the characterization of the fish pallium as a simpler neural structure, lacking the organizational sophistication necessary to support higher-order cognitive functions. Yet the apparent absence of topographic sensory maps in fishes may reflect the scarcity of functional studies rather than a true lack of spatial organization.

To address this issue, the present study used in vivo wide-field optical imaging to map sensory-evoked neural activity in the pallium of adult goldfish. This technique enables the simultaneous recording of activity patterns across the dorsal telencephalic surface with high spatial and temporal resolution, allowing for the precise delineation of sensory-responsive territories and their functional topography (Ferezou et al, 2009). We focused on non-olfactory modalities—somatosensory, auditory, gustatory, and visual—to determine (i) the number and location of sensory-responsive areas in the pallium, (ii) whether these areas are unimodal or polymodal, and (iii) whether they exhibit spatially organized sensory maps (e.g., somatotopy, tonotopy, gustotopy). By refining the functional parcellation of the teleost pallium, our findings provide new insight into its internal organization and contribute to ongoing debates on pallial homology.

Furthermore, considering that ray-finned fishes are the sister group of sarcopterygians, understanding the organization of the teleost pallium has important implications for reconstructing the evolution of forebrain organization across vertebrates, offering valuable clues about the ancestral condition of the vertebrate forebrain and the evolutionary origins of cortical circuits involved in perception, cognition, and emotion.

## RESULTS

The present study used in vivo wide-field voltage-sensitive dye imaging to map and characterize sensory-evoked neural activity across the dorsal surface of the goldfish telencephalic pallium (Figures 1A–C). This technique enables the generation of detailed spatiotemporal maps of pallial responses to somatosensory, auditory, gustatory, and visual stimulation, allowing the delineation of their topography, modality specificity, and functional organization. By aligning the boundaries of functional activation maps with external morphological landmarks and cytoarchitectural features, we established a refined anatomical–functional parcellation of the goldfish pallium. These analyses revealed spatially distinct activation domains within the Dm and Dl regions corresponding to different sensory modalities.

**Figure 1.**
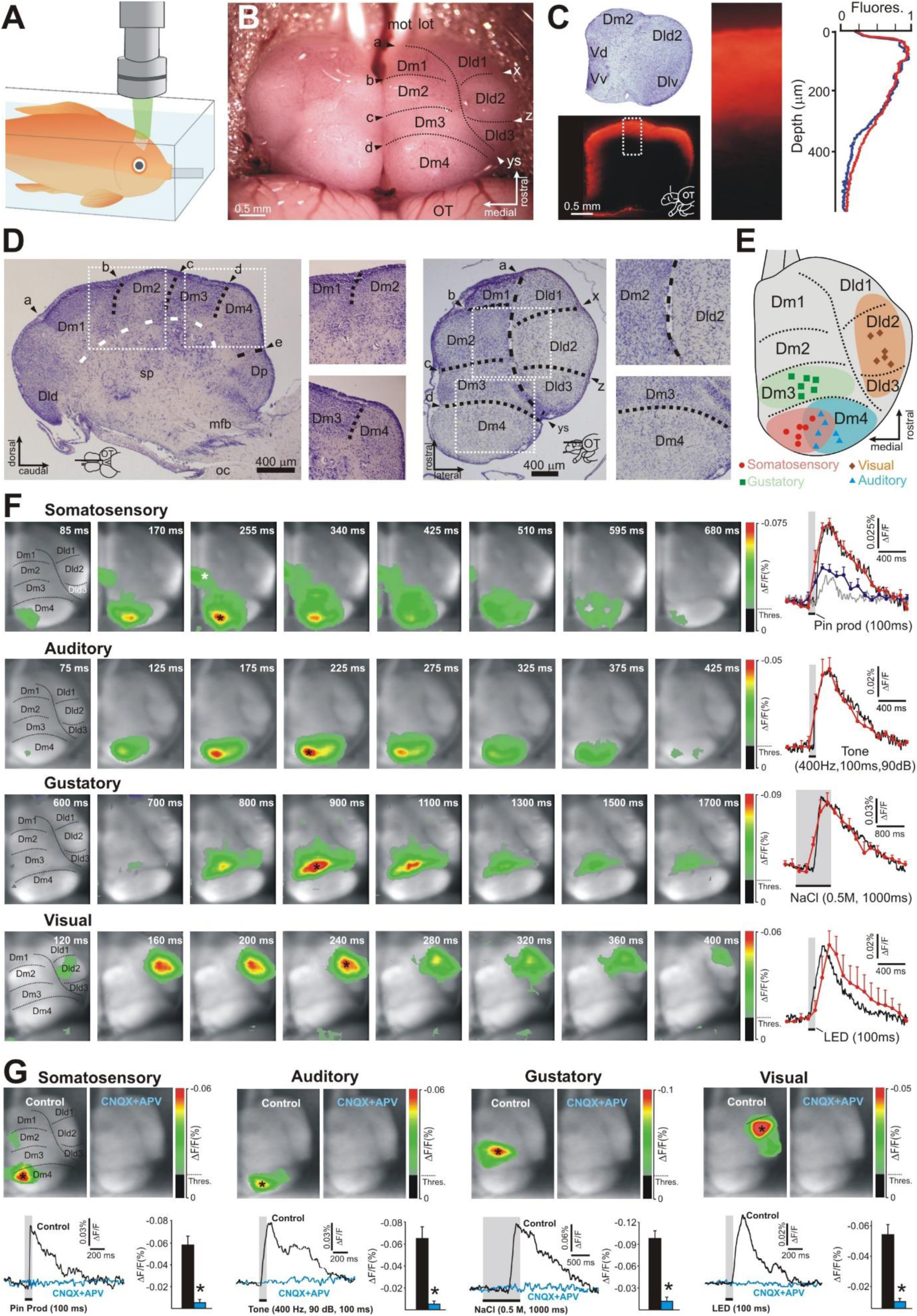
In vivo voltage-sensitive dye (VSD) imaging of sensory-evoked responses in the goldfish pallium. **A.** Schematic representation of the experimental setup, showing the fish in the recording chamber with a water flow tube inserted into the mouth to provide aerated water through the gills during the session, and the epifluorescence microscope positioned above for optical recording. **B**. Dorsal view of the goldfish brain showing the surgically exposed telencephalic hemispheres prepared for optical imaging. Pallial subdivisions identified in this study are outlined and labeled on the right hemisphere. **C**. Source of voltage-sensitive dye signals. Nissl- (upper left) and Di-2-ANEPEQ- (lower left) stained transverse sections through a telencephalic hemisphere showing the main pallial subregions and the depth of dye penetration. The curves (right) show relative fluorescence intensity as a function of depth along a radial strip of the pallium (indicated by the dashed white rectangle), corresponding to the slice shown on the left (blue line), and averaged across three subjects (red line). The VSD signal originates primarily within the upper 400 μm of the pallial surface. **D**. Cytoarchitectonic subdivision of the goldfish telencephalic pallium. Left: Nissl-stained sagittal section showing subdivisions of the Dm region. The white dashed line marks the pallium–subpallium boundary. Right: Nissl-stained horizontal section of a right telencephalic hemisphere. White squares indicate magnified views highlighting the Dm and Dl pallial regions and their respective subdivisions. **E**. Schematic representation of the location and extent of sensory areas identified in this study, shown in a dorsal view of the right telencephalic hemisphere. Each symbol marks the epicenter (defined as the 3 × 3 pixel region exhibiting maximal activation) of activity evoked by a specific sensory modality in a single subject. **F.** Spatiotemporal patterns of pallial activation in response to somatosensory, auditory, gustatory, and visual stimuli from representative subjects. Percent fractional fluorescence change at each pixel is color-coded according to the scale bar. Time elapsed since stimulus onset is indicated in each frame. Curves show the time course of the VSD signal measured from the region of maximal activation (3 × 3 pixel square; black asterisk) for the subject represented in each row (black line) and averaged across animals (red line; n = 6 per modality). Responses were characterized by an initial rapid decrease in fluorescence (membrane depolarization), followed by peak activity and a slower recovery phase (repolarization). Somatosensory stimulation evoked responses in two regions (Dm4 and Dm2; see text). For somatosensory responses, black and red curves correspond to Dm4 (single subject and group average, respectively), whereas grey and blue curves correspond to Dm2 (white asterisk; single subject and group average, respectively). **G.** Effects of ionotropic glutamate receptor blockade on sensory-evoked responses. Frames show responses evoked by each sensory modality before and after pharmacological blockade with CNQX + APV in a representative animal. For each modality, frames were obtained at the same post-stimulus time point (corresponding to peak response in the control condition). Curves represent time courses of VSD signals before (black) and after (blue) drug application, measured at the region of maximal activation in the control condition (black asterisk). Histograms show the effect of CNQX + APV averaged across animals (n = 4). Abbreviations: a–e, superficial indentations separating Dm subregions; Dld1–3, subdivisions 1–3 of the dorsal part of the lateral region of the area dorsalis; Dlv, ventral part of the lateral region of the area dorsalis; Dm1–4, subdivisions 1–4 of the medial region of the area dorsalis; Dp, posterior part of the area dorsalis; lot, mot, lateral and medial olfactory tracts; mfb, medial forebrain bundle; OT, optic tectum; sp, subpallium; Vd, Vv, dorsal and ventral regions of the area ventralis; x and z, superficial indentations separating Dld subregions; ys, ypsiloniformis sulcus.

### Distinct sensory areas are localized within anatomically defined subdivisions of the goldfish pallium

To determine the precise location and spatial relationships of the identified sensory areas, we integrated functional data with anatomical and histological landmarks. The goldfish telencephalic pallium is divided into medial (Dm) and lateral (Dl) regions, demarcated by the ypsiloniformis sulcus when viewed from the dorsal surface. The Dm region was further subdivided into four rostrocaudal subregions (Dm1–Dm4), while the dorsal portion of Dl (Dld) was subdivided into three rostrocaudal subdivisions (Dld1–Dld3). These subdivisions correspond to distinct bulges and valleculas readily discernible in dorsal view, exhibit characteristic cytoarchitectural patterns, and are separated by well-defined histological borders (Figures 1B,D). Together, these anatomical features provide a consistent framework for the functional mapping described below.

Sensory stimulation across modalities evoked activity in spatially segregated pallial domains (Figures 1E,F). Specifically, we identified a somatosensory domain located medially within Dm4, an auditory domain positioned more laterally within Dm4 and partially overlapping the somatosensory representation, a gustatory domain confined to Dm3, and a visual domain localized in Dld2. Together, these findings demonstrate the presence of modality-specific sensory territories distributed across anatomically defined pallial subdivisions, forming the basis for the topographic analyses presented in the following sections.

### Sensory-evoked responses are mediated by ionotropic glutamate receptors

To investigate the neurotransmitter mechanisms underlying sensory activity in the goldfish pallium, we pharmacologically blocked ionotropic glutamate receptors and quantified the resulting changes in neural responses. Baseline evoked activity to visual, auditory, somatosensory, and gustatory stimulation was first recorded, followed by topical application of a mixture of the AMPA/kainate receptor antagonist CNQX (10 µM) and the NMDA receptor antagonist DL-APV (50 µM) to the pallial surface for 45 minutes. After this treatment, sensory-evoked responses were markedly reduced across all modalities compared to baseline recordings (Figure 1G). Statistical analysis confirmed a significant reduction in response amplitudes following receptor blockade (all t₃ > 5.474, all P < 0.012; n = 4), demonstrating that ionotropic glutamatergic transmission is required for sensory activation of pallial circuits. This pharmacological manipulation indicates that the voltage-sensitive dye signals predominantly reflect excitatory synaptic population activity, thereby validating the functional mapping approach used in this study.

### Somatosensory-evoked activity in the pallium

A mild somatosensory stimulus (brief 100 ms touch at the base of the dorsal fin; n = 6 subjects) consistently evoked a response in the Dm4 subregion. Dm4 is the most caudal subdivision of Dm, separated from Dm3 at the midline by an indentation in the medial wall (labeled d in Figure 1B,D), from the posterior part of the area dorsalis (Dp) caudally by indentation e (Figure 1D), and from Dld by the ypsiloniformis sulcus.

Cytoarchitectonically, Dm4 contains scattered, medium-sized cells arranged in multiple layer-like structures, and its border with Dm3 is marked by an abrupt transition to the densely packed, darkly staining, granule-like small cells characteristic of Dm3 (Figure 1D).

The spatial extent and location of somatosensory responses were highly consistent across individuals (Figure 2A). The response was characterized by an initial rapid depolarization, followed by a peak of activity and a slower repolarization to baseline (Figures 1F and 2A). The earliest response appeared in medial Dm4 with a latency of 85.0 ± 7.2 ms, subsequently spreading laterally across Dm4, and reaching peak activity at 271.7 ± 18.2 ms (Figure 2A).

**Figure 2.**
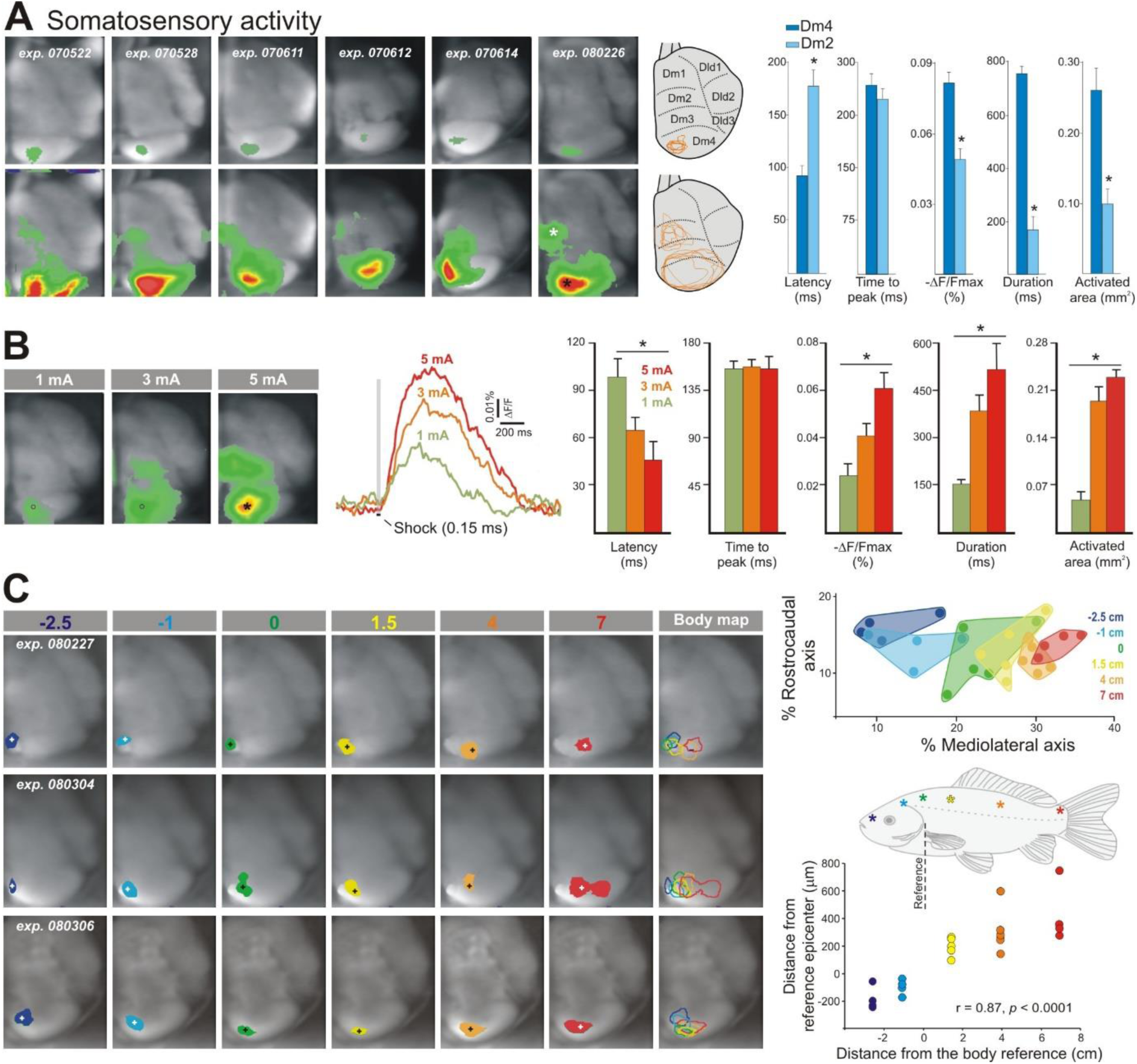
Somatosensory-evoked activity. **A.** Optical images of early (top; 25% of peak amplitude) and peak (bottom) somatosensory-evoked responses (n = 6). Schematic representations of the telencephalic surface on the right show superimposed contour plots of early and peak responses. Activated area outlines were aligned using the external borders of the telencephalon and the ypsiloniformis sulcus as landmarks. The relative location and extent of responsive areas were highly consistent across subjects. The histograms show quantitative parameters of the evoked responses in Dm4 and Dm2. The peak amplitude, activated area, and response duration were higher in Dm4 than in Dm2 (all t₅ > 3.952, all P < 0.011), whereas the response latency was shorter in Dm4 than in Dm2 (t₅ = −8.006, P < 0.001). **B.** Effects of electrical shock intensity on peak responses in a representative subject (left). Activity traces were measured from the regions marked by a black asterisk in the optical images (n = 4). Bar graphs show the effects of stimulus intensity on response parameters (right). Increasing stimulus intensity significantly increased peak amplitude (F_2,11_ = 8.704, P = 0.01), activated area (F_2,11_ = 38.576, P < 0.001), and depolarization duration (F_2,11_ = 6.870, P = 0.018), and significantly decreased onset latency (F_2,11_ = 5.640, P = 0.03). In contrast, stimulus intensity did not significantly affect time to peak (F_2,11_ = 0.012, P = 0.988). **C.** Optical images of peak responses evoked by mechanical stimulation at different body positions in three representative subjects (left panel). Numbers above each column indicate the six stimulated body positions (diagram of the fish body), measured in centimeters relative to the reference point (caudal tip of the operculum; position 0). Negative values indicate rostral positions. Images were thresholded at 80% of peak response and binarized using a distinct color code for each body position (core activation). The last column shows superimposed outlines of activated domains across positions. A color-coded somatotopic map (bottom right) was constructed based on the locations of response epicenters (3 × 3 pixel regions of maximal activation). Epicenter positions (plus symbols in the left panel) were normalized to the total length of the mediolateral and rostrocaudal axes of each telencephalon. The scatter plots show the correlation between epicenter position and stimulated body position along the mediolateral and rostrocaudal axes, respectively, with stimulation sites indicated on the goldfish body diagram.

Approximately 85 ms after onset of the Dm4 response, a second somatosensory domain was activated in Dm2 (white asterisk in Figure 2A). This secondary response was weaker and shorter in duration and was confined to a prominent bulge in Dm2 visible as an anatomical landmark (Figures 1B,D). The Dm2 subregion is delimited rostrally by indentation b (Figure 1B,D), separating it from Dm1, and caudally by indentation c (Figure 1B,D), marking its boundary with Dm3. Cytoarchitectonically, Dm2 is composed of scattered, medium-sized neurons that contrast with the densely packed small-cell populations characteristic of Dm1 and Dm3 (Figure 1D). Dm2 is separated from Dld by the ypsiloniformis sulcus.

To examine the functional specificity of Dm4 and Dm2 in somatosensory processing, we applied electrical shocks of increasing intensity to the skin (150 µs, 1–5 mA; n = 4 subjects; Figure 2B). Whereas Dm4 was activated in a graded manner at all stimulation intensities, Dm2 was recruited exclusively at higher intensities. These findings suggest that Dm2 represents a functionally distinct subregion preferentially engaged by high-intensity stimulation, potentially contributing to the encoding of stimulus salience or valence.

### Somatotopic organization in the somatosensory area

To determine whether the somatosensory domain in Dm4 encodes a spatial representation of the body surface, we applied mechanical stimulation at six standardized sites along the rostrocaudal axis of the fish body (n = 6 subjects; Figure 2C), using the caudal tip of the operculum as reference point (position = 0). At least three positions were tested per subject.

Topographic analysis was performed by thresholding activation maps at 80% of peak signal (core activation) and superimposing the resulting contour plots to compare response locations. Core activation zones for different body regions, although partially overlapping, systematically shifted along the mediolateral axis of Dm4 (Figure 2C, left panel).

Epicenters—defined as the 3 × 3 pixel region of maximal intensity—were likewise systematically aligned along the mediolateral axis (Figure 2C, upper right panel). Stimulation of rostral body sites (e.g., –2.5 cm from the reference) activated the most medial portions of Dm4, whereas progressively more caudal stimulation sites (e.g., +4 to +7 cm) shifted activation laterally. Quantitative analysis confirmed a strong linear correlation between epicenter position and stimulation site (r = 0.87, P = 0.0001; Figure 2C, bottom right panel).

Together, these findings demonstrate the presence of a robust somatotopic organization within Dm4 with a medial-to-lateral representation of the body axis, thereby providing the first direct evidence of spatial body mapping in the teleost pallium.

### Auditory-evoked activity in the pallium

Auditory stimulation (400 Hz, 90 dB, 100 ms; n = 6) evoked a consistent depolarization response restricted to Dm4 (Figures 1E,F and 3A). The Dm4 bulge (delimited rostrally by indentation d, caudally by the indentation e, and laterally by the ypsiloniformis sulcus; Figure 1D) serves as a reliable anatomical landmark for localizing the auditory pallial area, as the core activation rarely extended beyond its boundaries (Figure 3A).

**Figure 3.**
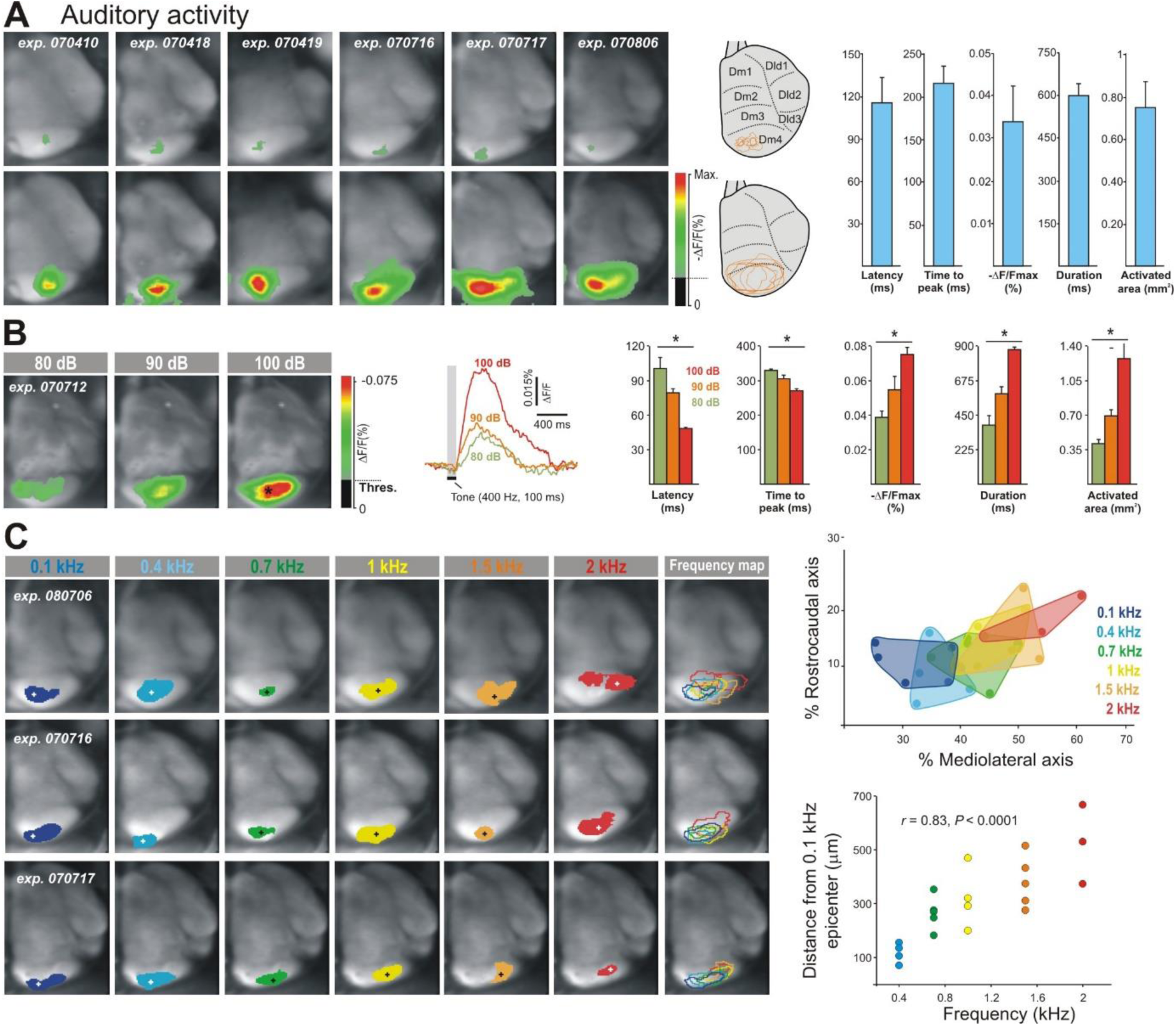
Auditory-evoked activity. **A**. Optical images of early (top; 25% of peak amplitude) and peak (bottom) auditory-evoked responses (n = 6). Schematic representations of the telencephalic surface on the right show superimposed contour plots of early and peak responses. Activated area outlines were aligned using the external contours of the telencephalon and the ypsiloniformis sulcus as anatomical landmarks. The spatial distribution of responsive areas was highly consistent across subjects. The histograms show quantitative parameters of the evoked responses. Abbreviations as in Figure 1. **B**. Effects of varying auditory stimulus intensity. Frames (right) show peak responses evoked by tones of different intensities. Curves illustrate the time course of the optical signal elicited by each stimulus, recorded at the site marked by the black square in the images (left). Histograms summarize the mean effects of stimulus intensity on response parameters (n = 4). **C**. Activity evoked by tones of six different frequencies (0.1–2 kHz; 100 ms; 90 dB) in three representative animals (left panel). Images show core activations (responses thresholded at 80% of peak amplitude and binarized using a distinct color code for each frequency). The last column shows superimposed outlines of activated domains across frequencies to illustrate their systematic spatial shift as a function of stimulus frequency. Upper right, color-coded tonotopic map reconstructed from epicenter locations (plus symbols in the left panel). Epicenter positions were normalized as percentages of the maximum length of the mediolateral and rostrocaudal telencephalic axes of each subject. Bottom right, scatter plot showing the correlation between stimulus frequency and epicenter position along the mediolateral axis, expressed as distance relative to the 0.1 kHz epicenter.

The auditory representation was positioned laterally relative to the Dm4 somatosensory representation and partially overlapped with it (Figures 1E–F). No clear cytoarchitectonic boundary was observed between the somatosensory and auditory representations within Dm4 (Figure 1D). Contour plots of the early response (25% of peak amplitude) and peak response showed consistent localization within Dm4 across animals (Figure 3A). The mean onset latency of auditory-evoked activity was 115.63 ± 17.89 ms. Over the subsequent milliseconds, the optical signal propagated along the mediolateral axis of Dm4 and peaked at 215.37 ± 19.41 ms.

The effects of varying sound intensity were examined (n = 4). Optical responses to pure tones (400 Hz, 100 ms) at different sound pressure levels (80, 90, and 100 dB) revealed significant modulation of multiple response parameters within Dm4 (Figure 3B). Specifically, increasing sound pressure level significantly decreased response latency (F₂,₁₁ = 53.622, P = 0.0001) and time to peak (F₂,₁₁ = 12.737, P = 0.002), while significantly increasing amplitude (F₂,₁₁ = 10.682, P = 0.004), duration (F₂,₁₁ = 25.818, P = 0.0001), and activated area (F₂,₁₁ = 15.774, P = 0.001).

### Tonotopic maps in the auditory area

We next examined whether the goldfish pallium contains a spatial representation of sound frequency. Animals (n = 6) were exposed to tones of different frequencies (0.1, 0.4, 0.7, 1, 1.5, and 2 kHz) with constant duration and intensity (100 ms, 90 dB). A minimum of three frequencies were tested per animal. Core activation domains and epicenters were defined as described for the somatosensory analysis. Although partially overlapping, activation domains systematically shifted along the mediolateral axis of Dm4 as stimulus frequency increased (Figure 3C, left panel). Similarly, epicenters displayed a consistent medial-to-lateral displacement with increasing frequency (Figure 3C, upper right panel). Low-frequency tones (0.1 kHz) produced epicenters in the most medial portion of Dm4, whereas high-frequency tones (2 kHz) produced epicenters located more laterally, with intermediate frequencies occupying intermediate positions. A significant linear correlation was observed between epicenter position (measured relative to the 0.1 kHz epicenter) and stimulus frequency (r = 0.83, P = 0.0001; Figure 3C, bottom right panel).

Together, these results demonstrate a mediolaterally organized tonotopic map within Dm4. Although the extent of the tonotopically responsive area varied slightly among individuals, the orderly frequency representation was highly consistent across subjects (Figure 3C).

### Gustatory-evoked activity in the pallium

Gustatory stimulation (intraoral 0.5 M NaCl, 1 s; n = 6) evoked a consistent response across animals that began in the caudal portion of Dm3, just rostral to the Dm4 bulge, and gradually expanded to encompass nearly the entire Dm3 region (Figures 1F and 4A). Dm3 is delimited rostrally from Dm2 by indentation c, caudally from Dm4 by indentation d, and laterally from Dld by the ypsiloniformis sulcus (Figure 1B,D).

**Figure 4.**
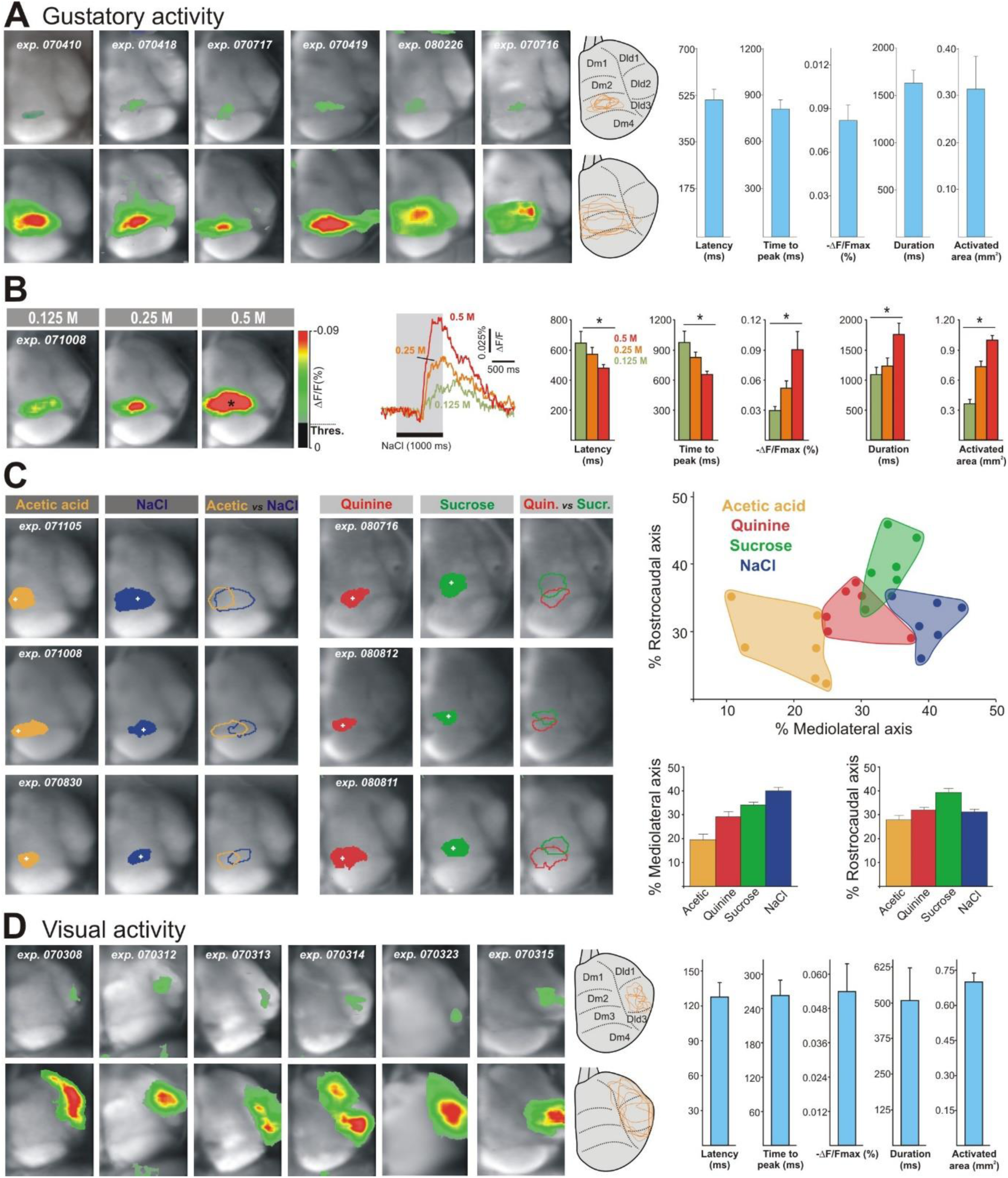
Gustatory-evoked (A-C) and visually-evoked (D) activity. **A**. Early (top panels) and peak (bottom panels) gustatory-evoked responses to NaCl solution (n = 6). Schematic representations of the telencephalic surface on the right show superimposed contour plots of early and peak responses, with outlines spatially aligned to the external contours of the telencephalon and the ypsiloniformis sulcus. Responsive areas exhibited a highly consistent spatial distribution across subjects. Histograms summarize the quantitative parameters of the evoked responses. Abbreviations as in Figure 1. **B**. Effects of varying tastant concentration. Frames show peak responses to different NaCl concentrations. Curves depict the time course of the optical signal evoked by each concentration, recorded at the sites marked by a black square in the images (left). Histograms summarize the mean effects of NaCl concentration on response parameters (n = 4). **C**. Optical images of peak responses evoked by four tastants (0.5 M NaCl vs 0.05 M acetic acid; 0.5 M sucrose vs 10⁻⁴ M quinine hydrochloride) in six representative animals (left panel). NaCl and acetic acid were tested in three animals, and sucrose and quinine hydrochloride in a separate group of three animals. Responses (core activations) were thresholded at 80% of peak amplitude and binarized using a distinct color code for each tastant. Superimposed outlines of domains activated by paired tastants in each animal are shown. Upper right, color-coded gustotopic map reconstructed as described in Figure 3C. Histograms show mean epicenter positions for each tastant, expressed as percentages of the maximum length of the mediolateral and rostrocaudal telencephalic axes (n = 6). Bottom right, scatter plots show the distribution of epicenter positions for each tastant along the mediolateral and rostrocaudal axes of Dm3. Linear correlation analyses were performed for each axis, analogous to the somatosensory and auditory analyses. **D**. Early (top panels) and peak (bottom panels) visually-evoked responses (n = 6; details as in Figure 2A). Schematic representations of the telencephalic surface show superimposed contour plots of early and peak responses. The spatial distribution of responsive areas was highly reproducible across subjects. The histograms present the quantitative parameters of the evoked responses. Abbreviations as in Figure 1.

Cytoarchitectonically, Dm3 is characterized by densely packed, darkly staining, granule-like small cells. This pattern contrasts with adjacent subregions: the Dm4–Dm3 border is marked by an abrupt transition from the scattered, medium-sized, layer-like cells typical of Dm4, whereas the Dm2–Dm3 border shows a shift from the more loosely distributed, medium-sized neurons characteristic of Dm2 (Figure 1D).

Gustatory-evoked responses exhibited a latency of 505.6 ± 40.8 ms, a duration of 1,635.0 ± 138.0 ms, and a time to peak of 810.7 ± 158.3 ms (Figure 4A). To determine whether these response dynamics were modulated by stimulus intensity, we examined the effect of tastant concentration on gustatory activity. Increasing NaCl concentrations (0.125, 0.25, and 0.5 M) produced systematic modulation of activity within Dm3 (Figure 4B), with significant increases in peak amplitude (F₂,₁₁ = 13.455, P = 0.002), response duration (F₂,₁₁ = 5.536, P = 0.027), and activated area (F₂,₁₁ = 14.848, P = 0.001), and significant decreases in response latency (F₂,₁₁ = 4.432, P = 0.046) and time to peak (F₂,₁₁ = 4.574, P = 0.043). These results indicate that the Dm3 gustatory area encodes stimulus intensity through graded changes in multiple response parameters.

### Spatial representation of taste stimuli in the gustatory area

To assess whether gustatory responses are spatially organized within Dm3, we recorded optical responses to intraoral delivery of different tastants (0.5 M NaCl, 0.5 M sucrose, 10⁻⁴ M quinine hydrochloride, and 0.05 M acetic acid; 1 s each; n = 12). Core activation domains and epicenters were defined as described above. Different tastants activated partially overlapping but spatially distinct domains within Dm3 (Figure 4C). In paired comparisons within the same fish (n = 4), the NaCl-evoked domain was positioned more laterally than that evoked by acetic acid. Similarly, sucrose and quinine hydrochloride activated distinct domains, with sucrose responses located more rostrally than quinine-evoked responses. Analysis of epicenter positions revealed significant differences along both the mediolateral (F₃,₂₃ = 22.231, P = 0.0001) and rostrocaudal (F₃,₂₃ = 9.428, P = 0.0001; Figure 4C) axes. Post hoc comparisons showed that acetic acid, quinine, and NaCl responses occupied distinct positions along the mediolateral axis, with acetic acid located medially, NaCl laterally, and quinine in intermediate positions (all P < 0.008). Epicenters for sucrose were significantly more rostral than those of the other tastants (all P < 0.023) and significantly more lateral than those evoked by acetic acid (P = 0.0001).

Together, these findings indicate a coarse but systematic spatial organization of gustatory representations within Dm3, with different tastants preferentially activating partially segregated subregions along both the mediolateral and rostrocaudal axes.

### Visually-evoked activity in the pallium

The visual stimulus (red light, 100 ms; n = 6) activated a well-defined domain within the Dld region (Dld2), with the core depolarization consistently located lateral to the ypsiloniformis sulcus (Figures 1E,F and 4D). Dld2 corresponds to a conspicuous bulge within Dld at the mid-hemispheric level and is separated rostrally from Dld1 by indentation x and caudally from Dld3 by indentation z (Figure 1B,D). Medially, Dld2 is delimited from Dm by the ypsiloniformis sulcus. Cytoarchitectonically, Dld2 is characterized by scattered small cells that stain more lightly than those of Dld1 and Dld3 (Figure 1D). Contour plots of the early and peak responses confirmed that the visual domain in Dld2 was highly reproducible across animals (Figure 4D).

## DISCUSSION

This study used in vivo wide-field voltage-sensitive dye imaging to map the sensory-evoked activity across the dorsal surface of the goldfish pallium. This technique allowed the identification of sensory-responsive areas for somatosensory, auditory, gustatory, and visual modalities with high spatial and temporal resolution. By aligning functional activity patterns with anatomical landmarks and cytoarchitectural features, we provide a refined parcellation of the goldfish pallium and new insights into its morpho-functional organization.

### Functional organization of the dorsomedial pallium (Dm)

Our results show that Dm is not a unitary division but a heterogeneous region comprising at least four rostrocaudally arranged subregions (Dm1, Dm2, Dm3, and Dm4). Within Dm, sensory responses were confined to Dm3 and Dm4, with somatosensory and auditory activity located in Dm4, and gustatory responses in Dm3. Somatosensory and auditory areas partially overlapped, while the gustatory region remained more segregated. These findings are consistent with previous electrophysiological and tract-tracing studies, which suggest partial segregation of sensory inputs to Dm (Echteler, 1985; Pretchl et al., 1999; Yamamoto & Ito, 2005).

Importantly, we report that sensory areas in Dm respond gradually to stimulus strength changes and are organized topographically. Somatosensory and auditory inputs form somatotopic and tonotopic maps in Dm4, while Dm3 shows a gustotopic organization. To our knowledge, this is the first demonstration of topographic sensory representations in the teleost pallium. The partial overlapping of these sensory representations in Dm3 and Dm4 could facilitate multisensory integration, supporting the possibility of unified perceptual experiences in fish. In contrast, Dm2 activates only when the strength of somatosensory stimulation increases to aversive levels.

This pattern of response suggests a functional dissociation: while Dm4 may encode sensory-perceptual components of the stimulus-such as modality, location, and intensity-Dm2 could be specifically involved in the representation of the affective or arousal-related aspects of nociceptive or aversive stimulation. Supporting this view, complementary studies from our lab have shown that Dm2 is a key center for emotional activation, Pavlovian conditioning, and place aversive learning in goldfish (Del Águila, 2024; Quintero, 2024), further reinforcing its role as an emotional processing center. Similar findings in zebrafish support a conserved function for this region across teleosts (Aoki et al., 2013; Lal et al., 2018).

Interestingly, an unexpected finding of the present study is that a large portion of the goldfish telencephalic pallium, namely Dm1, did not show sensory-evoked activity, indicating the presence of “sensory silent” area, likely involved in higher-order processing functions. In agreement with the present observations, previous connectional studies have shown a scarcity of diencephalic-driven sensory inputs to the rostral regions of the pallium compared to the caudal areas, together with their rich intrapallial connectivity (Northcutt, 2006, Yañez et al., 2021), which supports the idea that Dm1 may serve integrative functions.

In regard to the possible comparative identity of Dm some contentious hypotheses have been proposed. This controversy echoes the persistent debate regarding the identification of pallial homologies across non-avian reptiles, birds, and mammals (Finger et al, 2013; Jarvis et al., 2005; Medina & Abellán 2009; Puelles 2001; Wang et al., 2010). One view equates the sensory-recipient areas of Dm with the mammalian neocortex, based on topological position, segregated sensory domains, and similar molecular markers and cortical-like gene expression (Butler 2000; Northcutt 1981; Mueller et al., 2011; Thrin et al, 2025; Wullimann & Mueller, 2004; Yamamoto et al., 2007). However, this proposal is weakened by the fact that sensory inputs to the teleost pallium are routed via the preglomerular complex—a hypertrophied structure typical of teleosts derived from the midbrain and posterior diencephalon—rather than through thalamic relays (Bloch et al 2020; Ishikawa et al 2007; Northcutt, 2008; Yamamoto & Ito, 2005). Furthermore, unlike the mammalian neocortex, Dm lacks fully segregated unimodal areas (as shown here, sensory representations in Dm3 and Dm4 are only partially segregated and show substantial overlap across modalities), is not essential for stimulus discrimination (Broglio et al 2010; Martín-Monzón et al 2011; Ohnishi, 1997), is not involved in sensoriomotor functions (Quintero, 2024), maintains rich hypothalamic and visceral reciprocal connectivity (Kato et al.,2011; Northcutt, 2006), and is involved in emotional processing and activation (Amores, 2024; Lal et al 2018; Portavella et al 2004)-features that challenge the comparison with the neocortex.

An alternative hypothesis posits that Dm is homologous to the pallial amygdala (Braford, 1995; Broglio et al., 2005; Portavella et al., 2004; Salas et al., 2006). This view finds some support in its connectivity patterns, gene expression profiles, and its involvement in emotional learning and motivation (Ganz et al., 2014; Northcutt, 2006; Porter & Mueller, 2020; Wullimann & Mueller, 2004). However, it fails to account for the presence of partially segregated, modality-specific domains and topographic sensory maps in Dm3 and Dm4, features that are not characteristic of a purely amygdalar territory.

Thus, neither of the two prevailing homology hypotheses of Dm—whether equating it with the neocortex or with the pallial amygdala—adequately accounts for the diversity of structural, functional, and connectional features observed in this region. Each hypothesis captures only a limited subset of the available evidence, while failing to accommodate key findings from our study, most notably the presence of partially segregated and topographically organized sensory domains in Dm3 and Dm4, as well as the affective specialization of Dm2. To address these limitations, we propose the alternative, integrative hypothesis that Dm is neither homologous to the neocortex nor to the amygdala, but rather corresponds to a broader mammalian mesocortical system. Specifically, we propose that Dm is composed of functionally specialized subregions comparable to different components of the mammalian mesocortical (cortico-limbic) network. In particular, Dm3 and Dm4 exhibit an insular cortex-like organization, sharing key functional, such as modality-specific topographic maps for gustatory, somatosensory, and auditory inputs (Bechara & Damasio, 2005; Craig, 2009; Gogolla, 2017; Rolls, 2013). Unlike the amygdala, they respond to neutral stimuli (present results), and their lesion or stimulation does not directly affect emotional processing (Del Águila, 2024; Reiriz, 2017). In turn, Dm2 functionally parallels the anterior cingulate and medial prefrontal cortices through its involvement in emotional arousal and affective processing (Palomero-Galagher & Amunts, 2022; Rolls, 2019; Vogt 2005). This integrative hypothesis provides a more comprehensive account of the available evidence and helps to reconcile conflicting interpretations regarding the functional organization of Dm.

### Visual processing in Dld

The only pallial visual area was identified in Dld2, a subregion of the dorsolateral telencephalon. The visual area was confined to the Dld2 region, and its functionally defined boundaries closely matched well defined morphological and cytoarchitectural landmarks. In contrast, the adjacent regions Dld1 and Dld3 were unresponsive to primary sensory stimulation. These findings refine earlier functional and hodological studies and confirm that Dld2 constitutes the principal visual area in the teleost pallium (Bloch et al 2020; Hagio et al 2018; Ito & Vanegas 1983, 1984; Prechtl et al., 1998; Rakic et al., 1979; Saidel et al., 2001; Yamamoto & Ito 2008).

Several conflicting hypotheses regarding the homology of Dld have been proposed, echoing the long-standing debate about the evolutionary origins of the pallial visual projections in mammals and birds (Karten, 1991). One proposal equates Dld with neocortical visual areas, based on its sensory inputs and topological position within the pallium (Butler 2000; Trihn et al,2025; Wullimann & Mueller, 2004; Yamamoto et al 2007). However, as in Dm, visual information reaches Dld via the preglomerular complex rather than through thalamic relays, undermining its comparability to thalamorecipient neocortical areas (Bloch et al 2020; Northcutt, 2008; Yamamoto & Ito, 2005). An alternative hypothesis identifies Dl-including both Dld and Dlv-as homologous to the medial pallium (hippocampus) (Northcutt, 2006; Porter & Mueller, 2020), yet this view also encounters significant difficulties. While Dlv is involved in spatial memory and shares molecular markers with the hippocampus, Dld does not (Anneser et al 2024; Giassi et al, 2012; Hegarty et al 2024; Rodríguez et al., 2002). Lesions to Dld do not affect spatial learning, and it does not show plasticity-related changes after navigation tasks (Salas et al., 2006). Moreover, the presence of a specialized visual area is not a feature characteristic of the hippocampus. Taken together, this evidence challenges both the neocortical and the hippocampal homology hypothesis for Dld, and highlights the need for a revised framework that accounts for its distinctive structural and functional profile.

Based on these considerations, we propose the alternative hypothesis that Dld can be compared to mesocortical retrohippocampal regions of mammals-such as the posterior cingulate, retrosplenial, and entorhinal cortices-while Dlv may correspond to the hippocampus proper, including the subiculum, CA1-CA3 subfields, and dentate gyrus. Supporting this view, it has been reported that the intrinsic circuit architecture of Dld in gymnotiform fish closely resembles that of mammalian transitional cortices (Giassi et al., 2012). Further support for this interpretation comes from transcriptomic studies that identify conserved neuronal populations in Dld and Dlv of teleosts that mirror the retrosplenial–subicular–hippocampal axis in mammals (Hegarty et al., 2024; Pandey et al., 2023; Tibi et al., 2023). Specifically, we suggest that Dld2 may functionally correspond to the mammalian retrosplenial cortex (Mueller et al., 2011; Palomero-Gallagher & Amunts, 2022; Vogt, 2005), a region involved in higher-order visual processing, spatial integration, and contextual memory, rather than in primary sensory visual functions.

### Concluding remarks

The findings of the present study reveal that the teleost pallium possesses a more elaborate and functionally differentiated internal organization than previously acknowledged. The identification of discrete, modality-specific sensory areas exhibiting topographic order challenges the traditional view of a diffusely organized pallium in ray-finned fishes. Moreover, our results expose the limitations of existing homology hypotheses, which fail to account for the functional complexity of the teleost pallium and the co-occurrence of sensory, affective, and integrative domains within regions such as Dm and Dld. To address this gap, we propose a new interpretative framework in which these pallial territories are viewed as functionally comparable to components of the mammalian mesocortex, including insular, cingulate, and retrohippocampal cortices. This proposal integrates anatomical, functional, and connectional evidence into a coherent model for comparing pallial organization in teleost fishes with that of other vertebrates.

## METHODS

### Animal preparation

Goldfish (Carassius auratus), measuring 10–11 cm in standard length (snout to base of the caudal fin), were obtained from the vivarium of the University of Seville. Animals were anesthetized by immersion in a 1:20,000 solution of tricaine methanesulfonate (MS-222; Sigma-Aldrich; pH adjusted to 7.0–7.5) and placed in an experimental chamber equipped with an adjustable oral tube connected to a pump providing a continuous flow of aerated anesthetic solution through the gills. The anesthetic concentration was maintained constant throughout the surgical procedure. The dorsal skin and skull overlying the telencephalon were removed under visual guidance using a binocular microscope (SZ61, Olympus). The underlying intracranial fatty tissue was aspirated, and the tela choroidea was carefully removed to expose the dorsal telencephalic surface. Following surgery, the anesthetic was flushed out and replaced with fresh water. Recovery of an alert state was confirmed by the resumption of spontaneous breathing and eye and fin movements. Di-2-ANEPEQ (JPW 1114; Molecular Probes) was used as the voltage-sensitive dye due to its high water solubility, favorable diffusion properties, sensitivity to small voltage changes, high signal-to-noise ratio, and minimal photobleaching (Ferezou et al., 2009). A stock solution (0.5 mg/ml in distilled water) was diluted in goldfish Ringer solution (116 mM NaCl, 2.9 mM KCl, 1.8 mM CaCl₂, 5 mM HEPES; pH 7.2; Sigma-Aldrich) to a final concentration of 50 μg/ml. A volume of 100 μl was topically applied to the exposed telencephalon for 45 minutes. The tissue was then rinsed twice with Ringer solution to remove unbound dye and maintain moisture. For recordings, animals were immobilized by intraperitoneal injection of Flaxedil (5 μg/g body weight; gallamine triethiodide, Sigma-Aldrich) to minimize movement artifacts. All procedures complied with Directive 86/609/CEE of the European Community Council and Spanish legislation (R.D. 53/2013).

### Stimuli

Somatosensory, auditory, gustatory, and visual stimuli were applied. Mechanical stimulation consisted of a 100 ms touch delivered with a stainless steel pin-probe (1 mm diameter) attached to a solenoid controlled via an opto-coupled interface. Stimuli were delivered at six positions along the rostrocaudal axis of the left flank, approximately 5 mm above the lateral line. Pure tones (100 ms; 0.1–2 kHz; 80–100 dB; abrupt rise/fall) were generated using an auditory stimulator (LE-150, Letica Scientific Instruments) and delivered through a loudspeaker positioned ∼50 cm above and behind the head. Sound pressure levels were calibrated with a digital sound level meter. Taste stimuli were delivered via programmable syringe pumps (Aladdin; World Precision Instruments) at 0.5 ml/s for 1 s into the oral water flow through silicone tubing (4 mm diameter). Tastants included NaCl (0.125–0.5 M), acetic acid (0.05 M), quinine hydrochloride (10⁻⁴ M), and sucrose (0.5 M) prepared in distilled water (Sigma-Aldrich). Chamber water was continuously renewed to prevent chemical accumulation. To exclude olfactory contamination, gustatory experiments were performed in animals with sectioned olfactory tracts. Visual stimulation consisted of a 100 ms flash from a red LED (620 nm; 200 lux; Agilent Technologies) positioned 3 cm from the left eye. The LED was enclosed in an opaque latex tube to prevent stray illumination. The right eye was covered with opaque vinyl.

### Optical imaging and analysis

Optical recordings were obtained using a MiCAM01 system (Scimedia/Brain Vision; Tominaga et al., 2000). The epi-fluorescence microscope (THT; Scimedia) was mounted on a vibration isolation table. Excitation was provided by a 150 W tungsten halogen lamp (MHF-G150LR; Moritex) filtered at 530 ± 3 nm. Emitted fluorescence (>590 nm) was collected by a CCD camera (90 × 60 pixels; 2.9 × 2.1 mm sensor area). Using a 0.63× objective (PLAN APO; Leica Microsystems) and 1× projection lens, a 4.6 × 3.3 mm field of view was captured. The imaging field was centered on the dorsal surface of the right telencephalon. Images were acquired at 200 Hz (5 ms/frame). Acquisition began 400 ms after shutter opening to avoid shutter opening artifacts. Each trial lasted 1,700 ms (3,400 ms for gustatory experiments). A 300 ms prestimulus baseline was recorded. Sixteen trials were averaged to improve signal-to-noise ratio. Intertrial intervals were 30 s (60 s for gustatory trials). Images were processed using BV-Analyzer (Brain Vision). Signals were detrended to correct for bleaching, spatially averaged (5 × 5 pixel filter), and low-pass filtered. Fluorescence changes were expressed as percentage fractional change (%ΔF/F), calculated relative to the average of the first eight frames of the prestimulus baseline.

Pseudocolor maps were thresholded at 25% of full-scale signal to reduce background noise. Red indicated maximal depolarization (largest fluorescence decrease), yellow intermediate, and green minimal changes. Upward deflections in time-course traces correspond to depolarization. Response parameters included peak amplitude (mean of a 3 × 3 pixel region over maximal activation), latency (time to 25% of peak), time to peak, duration (time above 25% peak), and activated area (pixels above 25% peak at maximum response).

### Source of voltage-sensitive dye signals

To assess dye penetration depth, thin coronal sections were examined as described by Kleinfeld and Delaney (1996). After imaging, three animals were deeply anesthetized (1:5,000 MS-222) and perfused transcardially with 0.1 M PBS. Brains were removed, rapidly frozen, and sectioned coronally at 10 µm using a cryostat.

Sections were examined under an epifluorescence microscope (Axioskop 2; Carl Zeiss). Fluorescence intensity profiles indicated maximal labeling within the first 200 µm below the surface (Figure 1C). At 300 µm fluorescence was ∼50% of maximum, and at 400 µm it fell below 25%. Pharmacological blockade of ionotropic glutamate receptors NMDA receptor antagonist DL-APV (50 µM) and AMPA receptor antagonist CNQX (10 µM) were dissolved in teleost Ringer solution and applied directly to the telencephalic surface during blockade experiments.

### Nissl staining

To define anatomical boundaries, brains were perfused with PBS followed by fixative (methanol:acetone:water, 2:2:1). Tissue was post-fixed, paraffin-embedded, and sectioned (20 μm; Leica RM2125RT) in coronal, horizontal, or sagittal planes. Sections were deparaffinized, hydrated, and stained with cresyl violet (0.5%) for histological analysis.

### Statistical analysis

Data are presented as mean ± SEM. Statistical analyses were performed using Student’s t-test or repeated measures ANOVA (IBM SPSS). Significance was set at P < 0.05.

## STATEMENTS & DECLARATIONS

### Author Contributions

All authors contributed to the study conception and design, material preparation, data collection and analysis. The manuscript was written by all the authors. All authors read and approved the final manuscript.

### Data availability

Data will be made available on request.

## REFERENCES

Amores L (2024). Telencephalic pallium and conditioned taste aversion in goldfish (Carassius auratus). Unpublished Doctoral thesis. University of Sevilla. https://hdl.handle.net/11441/160797

Anneser L, Satou C, Hotz H & Friedrich R (2024). Molecular organization of neuronal cell types and neuromodulatory systems in the zebrafish telencephalon. Current Biology 34, 2, 298–312.e4, 10.1016/j.cub.2023.12.003

Aoki T, Kinoshita M, Aoki R, Agetsuma M, Aizawa H, Yamazaki M, Takahoko M, Amo R, Arata A, Higashijima S, Tsuboi T & Okamoto H (2013). Imaging of Neural Ensemble for the Retrieval of a Learned Behavioral Program. Neuron. 2013 Jun 5;78(5):881–94. 10.1016/j.neuron.2013.04.009

Bechara A & Damasio A (2005). The somatic marker hypothesis: A neural theory of economic decision. Games and Economic Behavior 52: 336–372. 10.1016/j.geb.2004.06.010

Bloch S, Hagio H, Thomas M, Heuzé A, Hermel JM, Lasserre E, Colin I, Saka K, Affaticati P, Jenett A, Kawakami K & Yamamoto N (2020). Non-thalamic origin of zebrafish sensory nuclei implies convergent evolution of visual pathways in amniotes and teleosts. eLife 9:e54945. DOI: 10.7554/eLife.54945

Braford M R (1995). Comparative aspects of forebrain organization in the ray finned fishes: touchstones or not? Brain, Behavior and Evolution, 46(4-5), 259–274. 10.1159/000113278

Broglio C, Gómez A, Durán E, Ocaña FM, Jiménez-Moya F, Rodríguez F & Salas C (2005). Hallmarks of a common forebrain vertebrate plan: specialized pallial areas for spatial, temporal and emotional memory in actinopterygian fish. Brain Research Bulletin 66:277–281. 10.1016/j.brainresbull.2005.03.021

Broglio C, Rodríguez F, Gómez A, Arias J L & Salas C (2010). Selective involvement of the goldfish lateral pallium in spatial memory. Behavioural Brain Research, 210(2), 191–201. 10.1016/J.BBR.2010.02.031

Butler AB (2000). Topography and topology of the teleost telencephalon: a paradox resolved. Neuroscience Letters 293:95–98. 10.1016/s0304-3940(00)01497-x

Craig A (2009). How do you feel — now? The anterior insula and human awareness. Nature Reviews in Neuroscience 10, 59–70. 10.1038/nrn2555

Damasio A & Carvalho G (2013). The nature of feelings: evolutionary and neurobiological origins. Nature Reviews in Neuroscience 14, 143–152. 10.1038/nrn3403

Del Águila T (2024). Telencephalic mechanisms underlying the nociceptive, affective, and cognitive components of pain in teleosts: a model for research on cortical pain mechanisms and their evolution in vertebrates. Unpublished doctoral thesis. University of Sevilla. https://hdl.handle.net/11441/161793

Echteler S M (1985). Organization of central auditory pathways in a teleost fish, Cyprinus carpio. Journal of Comparative Neurology, A, 156, 267–280. 10.1007/BF00610868.

Feinberg TE & Mallat JM (2016). The ancient origins of consciousness. How the brain created experience. The MIT Press. Cambridge, Massachusetts. London, England.

Ferezou I, Matyas F & Petersen CCH (2009). Imaging the Brain in Action: Real-Time Voltage- Sensitive Dye Imaging of Sensorimotor Cortex of Awake Behaving Mice. In Frostig R (ed); In Vivo Optical Imaging of Brain Function. 2nd edition. Boca Raton (FL): CRC Press/Taylor & Francis. Chapter 6. PMID: 26844323

Finger TE, Yamamoto N, Karten HJ & Hof PR (2013). Evolution of the Forebrain — Revisiting the Pallium. Journal of Comparative Neurology, 521: 3601–3603. 10.1002/cne.23444

Folgueira M & Clarke J D W (2024). Telencephalic eversion in embryos and early larvae of four teleost species. Evolution & Development, 26, e12474. 10.1111/ede.12474

Furlan G, Cuccioli V, Vuillemin N, Dirian L, Muntasell AJ, Coolen M, Dray N, Bedu S, Houart C, Beaurepaire E, Foucher I & Bally-Cuif L (2017). Life-Long Neurogenic Activity of Individual Neural Stem Cells and Continuous Growth Establish an Outside-In Architecture in the Teleost Pallium. Current Biology 6; 27 (21):3288–3301.e3. 10.1016/j.cub.2017.09.052

Ganz J, Brand M, Kroehne V, Freudenreich D, Machate A, Geffarth M, Braasch I & Kaslin J (2014) Subdivisions of the adult zebrafish pallium based on molecular marker analysis. F1000Research 3. 10.12688/F1000RESEARCH.5595.2

Giassi A C C, Ellis W & Maler L (2012). Organization of the gymnotiform fish pallium in relation to learning and memory: III. Intrinsic connections. Journal of Comparative Neurology, 520: 3369–3394. 10.1002/cne.23108

Gogolla N (2017). The insular cortex. Current Biology Jun 19;27 (12): R580–R586. 10.1016/j.cub.2017.05.010

Graziano MSA (2019). Rethinking Consciousness: A Scientific Theory of Subjective Experience. W.W. Norton

Hagio H, Sato M & Yamamoto N (2018). An ascending visual pathway to the dorsal telencephalon through the optic tectum and nucleus prethalamicus in the yellowfin goby Acanthogobius flavimanus (Temminck & Schlegel, 1845). Journal of Comparative Neurology 526:1733–1746. 10.1002/cne.24444.

Hegarty BE, Gruenhagen GW, Johnson ZV, Baker CM, Streelman JT (2024). Spatially resolved cell atlas of the teleost telencephalon and deep homology of the vertebrate forebrain Communications Biology 7:612. 10.1038/s42003-024-06315-1

Ishikawa Y, Yamamoto N, Yoshimoto M, Yasuda T, Maruyama K, Kage T, Takeda H & Ito H. (2007). Developmental origin of diencephalic sensory relay nuclei in teleosts. Brain, Behavior and Evolution, 69, 87–95. 10.1159/000095197

Ito H & Vanegas H (1983). Cytoarchitecture and ultrastructure of nucleus prethalamicus, with special reference to degenerating afferents from optic tectum and telencephalon, in a teleost (Holocentrus ascensionis). Journal of Comparative Neurology, 221, 401–415. 10.1002/cne.902210404

Ito H & Vanegas H (1984). Visual receptive thalamopetal neurons in the optic tectum of teleosts (Holocentridae). Brain Research, 290,201–210. 10.1016/0006-8993(84)90938-7.

Jarvis E, Güntürkün O, Bruce L, Csillag A, Karten H, Kuenzel W, Medina L, Paxinos G, Perkel DJ, Shimizu T, Striedter G, Wild JM, Ball GF, Dugas-Ford J, Durand SE, Hough GE, Husband S, Kubikova L, Lee DW, Mello CV, Powers A, Siang C, Smulders TV, Wada K, White SA, Yamamoto K, Yu J, Reiner A & Butler AB (2005). Avian brains and a new understanding of vertebrate brain evolution. Nature Reviews in Neuroscience 6, 151–159. 10.1038/nrn1606

Kato T, Yamada Y & Yamamoto N (2011), General visceral and gustatory connections of the posterior thalamic nucleus of goldfish. Journal of Comparative Neurology, 519: 3102–3123. 10.1002/cne.22669

Kleinfeld D & Delaney KR (1996). Distributed representation of vibrissa movement in the upper layers of somatosensory cortex revealed with voltage-sensitive dyes. Journal of Comparative Neurology 375(1):89–108. 10.1002/(SICI)1096-9861(19961104)375:1<89::AID-CNE6>3.0.CO;2-K

al P, Tanabe H, Suster ML, Ailani D, Kotani Y, Muto A, Itoh M, Iwasaki M. Wada H, Yaksi E & Kawakami K (2018). Identification of a neuronal population in the telencephalon essential for fear conditioning in zebrafish. BMC Biology (2018) 16:45. 10.1186/s12915-018-0502-y

LeDoux JE (2021). As soon as there was life, there was danger: the deep history of survival behaviours and the shallower history of consciousness. Philosophical Transactions Royal Society B 377: 20210292.10.1098/rstb.2021.0292

Martín I, Gómez A, Salas C, Puerto A, Rodríguez F (2011) Dorsomedial pallium lesions impair taste aversion learning in goldfish. Neurobiology of Learning & Memory 96:297–305. 10.1016/j.nlm.2011.06.003

Medina L & Abellán A (2009). Development and evolution of the pallium. Seminars in Cell & Developmental Biology 20: 698–711. 10.1016/j.semcdb.2009.04.008

Mueller T, Dong Z, Berberoglu MA & Guo S.(2011). The dorsal pallium in zebrafish, Danio rerio (Cyprinidae, Teleostei). Brain Research 1381:95 –105. 10.1016/j.brainres.2010.12.089

Nieuwenhuys R (2011). The development and general morphology of the telencephalon of actinopterygian fishes: synopsis, documentation and commentary. Brain Structure & Function 215:141–157. 10.1007/s00429-010-0285-6

Nieuwenhuys R, Donkelaar HJ & Nicholson C (1998). The central nervous system of vertebrates. Springer.

Northcutt, R. G. (1981). Evolution of the telencephalon in nonmammals. Annual Review of Neuroscience, 4, 301–50. 10.1146/annurev.ne.04.030181.001505

Northcutt RG (2006) Connections of the lateral and medial divisions of the goldfish telencephalic pallium. Journal of Comparative Neurology 494:903–943. 10.1002/cne.20853

Northcutt RG (2008) Forebrain evolution in bony fishes. Brain Research Bulletin 75:191–205. 10.1016/j.brainresbull.2007.10.058.

Ohnishi K (1997). Effects of telencephalic ablation on short-term memory and attention in goldfish. Behavioural Brain Research 86(2):191–9. 10.1016/s0166-4328(96)02265-6.

Palomero-Gallagher N & Amunts K (2022). A short review on emotion processing_ a lateralized network of neuronal networks. Brain Structure and Function 227:673–684 10.1007/s00429-021-02331-7

Pandey S, Moyer AJ & Thyme SB (2023) A single-cell transcriptome atlas of the maturing zebrafish telencephalon. Genome Research Apr;33(4):658–671. 10.1101/gr.277278.122.

Portavella M, Torres B & Salas C (2004) Avoidance response in goldfish: emotional and temporal involvement of medial and lateral telencephalic pallium. Journal of Neuroscience 24:2335–2342. 10.1523/JNEUROSCI.4930-03.2004

Porter BA & Mueller T (2020). The zebrafish amygdaloid complex - functional ground plan, molecular delineation, and everted topology. Frontiers in Neuroscience, 14, 608. 10.3389/fnins.2020.00608

Prechtl JC, von der Emde G, Wolfart J, Karamürsel S, Akoev GN, Andrianov YN & Bullock TH (1998). Sensory processing in the pallium of a mormyrid fish. Journal of Neuroscience, 18, 7381–7393. 10.1523/JNEUROSCI.18-18-07381.1998

Puelles L (2001). Thoughts on the development, structure and evolution of the mammalian and avian telencephalic pallium. Philosophical Transactions Royal Soc Lond B Biol Sci 2001;356:1583–98. 10.1098/rstb.2001.0973

Quintero, B. (2024). The role of the telencephalic pallium in teleost fish in motivational and emotional processes. Unpublished doctoral Thesis. University of Sevilla. https://hdl.handle.net/11441/160845

Rakic L, Belekhova MG & Konevi D (1979). Visual projections in the telencephalon and diencephalon of the teleost Serranus scriva. Zhurnal Evolyutsionnoi Biokhimii i Fiziologii, 15, 357–366.

Reiriz M (2017). Functional and anatomical characterization of the teleost telencephalic pallium areas involved in emotional processing. Unpublished Doctoral Thesis. University of Sevilla. http://hdl.handle.net/11441/61895

Rodríguez F, López JC, Vargas JP, Gómez Y, Broglio C, Salas C (2002) Conservation of spatial memory function in the pallial forebrain of reptiles and ray-finned fishes. Journal of Neuroscience 22:2894–2903. 10.1523/jneurosci.22-07-02894.2002

Rolls ET (2013). Limbic systems for emotion and for memory, but no single limbic system. Cortex 62: 119–157.10.1016/j.cortex.2013.12.005

Rolls ET (2019). The cingulate cortex and limbic systems for action, emotion, and memory. In: Vogt BA (Ed) Cingulate Cortex. Handbook of Clinical Neurology 3rd Series. Elsevier.

Saidel WM, Marquez-Houston K & Butler AB (2001). Identification of visual pallial telencephalon in the goldfish, Carassius auratus: a combined cytochrome oxidase and electrophysiological study. Brain Research, 919: 82–93. 10.1016/s0006-8993(01)03001-3

Salas C, Broglio C, Durán E, Gómez A, Ocaña FM, Jiménez-Moya F & Rodríguez F. (2006). Neuropsychology of Learning and Memory in Teleost Fish. Zebrafish, 3(2), 157–171. 10.1089/zeb.2006.3.157

Striedter GF & Northcutt RG (2020). Brains through time. A natural history of vertebrates. Oxford. University Press.

Striedter GF & Northcutt RG (2021). The Independent Evolution of Dorsal Pallia in Multiple Vertebrate Lineages. Brain, Behavior and Evolution, 96(4–6), 200–211. 10.1159/000516563

Tibi M, Hayun SB, Hochgerner H, Lin Z, Givon S, Ophir O, Shay T, Mueller T, Segev R & Zeisel A (2023). A telencephalon cell type atlas for goldfish reveals diversity in the evolution of spatial structure and cell types. Science Advances. 2023 Nov 3;9(44):eadh7693. 10.1126/sciadv.adh7693

Trinh AT, Ostenrath AM, Castillo-Berges I, Kraus S, Cachin F, Serneels B, Kawakami K & Yaksi E (2025). Hierarchical processing of sensory information across topographically organized thalamocortical-like circuits in the zebrafish brain. bioRxiv September 16, 2025 10.1101/2025.09.15.675867

Vogt BA (2005). Pain and emotion interactions in subregions of the cingulate gyrus. Nature Reviews Neuroscience, 6, 533e544. 10.1038/nrn1704

Wang Y, Brzozowska-Prechtl A & Karten HJ (2010). Laminar and columnar auditory cortex in avian brain. PNAS 107(28):12676–81. doi:10.1073/pnas.1006645107.

Wullimann MF & Mueller T (2004) Teleostean and mammalian forebrains contrasted: evidence from genes to behavior. Journal of Comparative Neurology 475:143–162. 10.1002/cne.20183

Wullimann MF & Vernier P (2009) Evolution of the Telencephalon in Anamniotes. In.: Marc D. Binder, Nobutaka Hirokawa and Uwe Windhorst (eds). Encyclopedia of Neuroscience Springer-Verlag GmbH Berlin Heidelberg 2009 https://10.1007/978-3-540-29678-2_3151

Yamamoto N, Ishikawa Y, Yoshimoto M, Xue HG, Bahaxar N, Sawai N, Yang CY, Ozawa H & Ito H. (2007). A new interpretation on the homology of the teleostean telencephalon based on hodology and a new eversion model. Brain, Behavior and Evolution, 69(2), 96–104. 10.1159/000095198

Yamamoto N & Ito H (2005). Fiber connections of the anterior preglomerular nucleus in cyprinids with notes on telencephalic connections of the preglomerular complex. Journal of Comparative Neurology, 491, 212–233. 10.1002/cne.20681

Yamamoto N & Ito H (2008). Visual, lateral line, and auditory ascending pathways to the dorsal telencephalic area through the rostrolateral region of the lateral preglomerular nucleus in cyprinids. Journal of Comparative Neurology, 508, 615–647. 10.1002/cne.21717

Yáñez J, Folgueira M, Lamas I & Anadón R (2022). The organization of the zebrafish pallium from a hodological perspective. Journal of Comparative Neurology, 530(8), 1164–1194. 10.1002/cne.25268

